# Data driven analysis reveals shared transcriptome response and immune cell composition across aetiologies of critical illness

**DOI:** 10.1101/525865

**Authors:** Zsolt Zador, Alexander Landry, Michael Balas, John C. Marshall, Michael D. Cusimano

## Abstract

Sepsis and trauma are frequent and challenging health problems in critical care. The diversity of patient response to these conditions complicates both disease management and outcome prediction. Whole blood transcriptomics allows the analysis of response in the critically ill at a molecular level. Prior results in this field demonstrate robust and diverse genomic response in the acute phase and others have shown shared biological mechanisms across wide disease aetiologies. We hypothesize that specific biological mechanisms, particularly those related to immune processes, are shared between sepsis and trauma cohorts. These may serve as universal markers for patients vulnerable to a complicated clinical course and/or mortality. We present a systems level analysis of gene expression for a total of 317 patients with abdominal sepsis (51), pulmonary sepsis (108) or trauma (158) and compare them to healthy controls (68). Our results confirm that immune processes are shared across disease aetiologies in critical illnesses. We identify two consistent and distinct subgroups of critical illness: 1) increased dendritic cell and CD4 T helper fractions but suppressed neutrophils and 2) high neutrophils and otherwise suppressed leukocyte fractions. These subgroups validate in an independent cohort of 181 paediatric patients suffering from septic shock of diverse aetiologies. Furthermore, we found immune and inflammatory processes derived from gene co-expression networks were downregulated in subgroup 1. This paralleled prior results which find similar leukocyte configuration by deconvolving whole blood transcriptomics, and this leukocyte configuration associates with greater susceptibility to multi organ failure. We are the first to identify a patient subgroup with a preserved leukocyte configuration across aetiologies of critical illness, which may serve as a universal predictor of complicated clinical course/poor outcome.

## Introduction

Host responses to critical illness such as infection or trauma are highly variable between patients (1–3). Specific examples of this complexity include the reprioritization patterns in gene expression (2,4,5) and the coexistence of proinflammatory and immunosuppressed states (1–3,6,7) in critically ill patients. Understanding this diversity in systemic response is highly important to treatment planning and recovery prediction. Analysis of whole blood transcriptomics (WBT) may facilitate this by identifying individual patient response to a variety of stresses leading to marker discovery and improved outcome prediction. It can also give valuable insight into biological processes and mechanisms of disease. WBT involves rapid isolation of total blood leukocytes followed by purification of total RNA and high throughput quantification of gene expression levels (1,2). WBT analysis has been applied to predict outcome in trauma (5), identify clinically relevant subgroups in sepsis (6) and explain diverse response to vaccination (8). Physiological processes such as aging have also been associated with a transcriptomics pattern to establish the “molecular age” of patients (9).

“Systems level analysis” of transcriptomics data can associate gene expression patterns with molecular processes and ultimately biological/clinical phenotypes. Two well-established and widely used methods in systems biology are gene expression networks (10,11) and the deconvolution of cellular fractions in bulk tissue (12). Gene expression networks are able to identify mechanisms relevant to physiological and pathological states by detecting clusters of highly co-expressed transcripts (13–15). An additional advantage of this approach is the ability to capture low and additive gene expression signals which may otherwise be overlooked by conventional differential gene expression analysis. This network-based technique has been successfully applied in exploring complex biological phenotypes like Huntington’s disease (16), peripheral nerve regeneration (13), malignant primary brain tumors (17) or weight gain (11). Deconvolution of bulk transcriptomics data uses cell-specific markers to estimate the fraction of corresponding cellular types present in a tissue sample (12). This technique has provided insight into the immune landscape of cancer (18), the tumor microenvironment in glioblastoma multiforme (19) and the immune cell composition of sera in a variety of diseases (20).

Studies have shown shared transcriptomics responses across different aetiologies of sepsis (6) and a large scale comparison of different diseases also suggested substantial overlap in WBT signal (4). However, the characteristics of these shared mechanisms and their relevance remains less explored. We firstly hypothesize that transcriptomics response to a variety of critical illnesses organizes into biological processes and an immune profile which are shared across aetiologies. We further hypothesize responses that are more predictive of good outcome than others. In the first part of our study we applied gene co-expression networks to open source transcriptomics data (2,21) and explore biological mechanisms in critical illness: abdominal sepsis, pulmonary sepsis and trauma. Our research is novel in the following: 1) contemporary studies largely rely on differential gene expression analysis to explore underlying mechanisms in critical illness, while our study uses co-expression network to identify relevant genes. This approach is rooted in biological reality, and may capture additive biological signal which could otherwise be overlooked by conventional techniques; 2) Our study is the first to explore the heterogeneity of patient response across critical illness with different aetiologies (sepsis due to community acquired pneumonia/abdominal source and trauma). We confirm that immune processes are shared across these conditions and we further explore the immune cell composition to identify consistently homogenous patient subgroups in critical illness.

## Methods

### Database

We used open source datasets from the open source genetic repository Gene Expression Omnibus (GEO) (22): GSE11375 (5) for trauma and GSE65682 (MARS Consortium) for abdominal (23) and pulmonary (24) sepsis. Demographics for the cohort are summarized in Table 1. For verification of our results we use a paediatric cohort of 181 patients (GSE66099) suffering from septic shock (25).

**Table 1:** Study demographics. A: control, B: trauma, C: CAP, D: abdominal sepsis, E: Septic shock (unknown aetiology) *Of available data. ^#^Validation cohort

| GEO entry | N | Mean age (SD) | Sex (M:F) | Patient distribution |  |  |  |  | Mortality (%) |
| --- | --- | --- | --- | --- | --- | --- | --- | --- | --- |
|  |  |  |  | A | B | C | D | E |  |
| GSE11375 | 184 | 33.1 (11.1) | 115:69 | 26 | 158 | 0 | 0 | 0 | 7/184 (3.8) |
| GSE65682 | 201 | 58.1 (17.3) | 114:87 | 42 | 0 | 108 | 51 | 0 | 31/201 (15.4) |
| GSE66099<sup>#</sup> | 228 | N/A | N/A | 47 | 0 | 0 | 0 | 181 | N/A |
| Overall* | 613 | 46.2 (19.2) | 229:156 | 68 | 158 | 108 | 51 | 181 | 38/385 (9.9) |

### Systems level analysis of whole blood gene expression data

The trauma cohort consisted of 158 adult patients under the age of 55 who suffered severe blunt trauma. Blood samples were taken within 12 hours of the injury and processed for gene expression analysis as described by Xiao et al (2). The cohort of patients suffering from sepsis (GSE65682) was divided into two groups based on the admitting: sepsis due to community acquired pneumonia (CAP) and abdominal sepsis (AS). Blood samples were taken within 24 hours of admission to critical care (23,24). To demonstrate the generalizability of our findings, we chose a large dataset of 276 paediatric patients, the Genomics of Pediatric SIRS and Septic Shock Investigators (GPSSSI)(25). In our verification subgroup, we had 181 unique patients who have been admitted with a diagnosis of septic shock from a variety of infectious sources and blood sample drawn on the first day of admission.

Gene co-expression patterns were identified using the well-established “Weighted Gene Correlation Network Analysis” (WGCNA) to detect “modules” (clusters) of strongly co-expressed genes (10). Per these previously described techniques, we first computed an “adjacency matrix”, a symmetrical matrix carrying soft-thresholded Pearson correlations between each gene pair. Computation of the similarity measures is detailed by Langfelder et al (10). This was then converted into a biologically-inspired topological overlap map (TOM), wherein pairwise gene similarities were derived from comparing their connectivity profiles (26). Hierarchical clustering with subsequent “dynamic” tree-cut (27) ultimately yielded gene modules, whose biological functions were annotated with the *Database for Annotation, Visualization and Integrated Discovery* (DAVID) (28), an open-source bioinformatics resource. Additionally, representative module “meta-genes” for each sample were computed as the first principal component of their constituent genes’ expression values, a well-established approach. We demonstrated associations between gene modules (thereby biological mechanisms) and external traits by correlating module meta genes with trait labels (for example as healthy vs septic) as described previously (10). Shared mechanisms as annotated by DAVID were identified based on ontological similarities using the GoSemSim package (29). This method determines similarities between Gene Ontology (GO) terms based on graph-based measures, wherein mechanisms are associated based on evolutionary and biological relationships and similarity is computed based on graph topology structure. Methods are described further in ref (29).

### Deconvolution of leukocyte cell factions

We used the well-established technique of CIBERSORT (12) to estimate the immune cell composition of whole blood. This method relies upon a previously derived reference gene expression matrix generated from a collection of pure lymphocyte fractions. The signature matrix was truncated to the genes detectable in all 3 cohorts (514 genes out of 547 genes in the original signature matrix). Within this signature matrix, each cell type is assigned a series of markers, which are then incorporated into nu-support vector regression to derive cell fractions. The method is further described in ref (12). The CIBERSORT technique is implemented from R package *epiDISH*.

### Patient subgroup discovery and validation

We used k-means clustering to identify patient subclasses based on immune cell landscape. This is a widely used technique in computational biology which initiates with a specified number of clusters (*k*) and assigns data points to each cluster based on a distance measure (30). During an iterative process, the position of the cluster centres in the data space are adjusted based on the cluster memberships as points are being added. Data points are reassigned as the cluster centre is being adjusted. The process continues until there is further adjustment required to the cluster centres (clusters are deemed stable). The optimal class numbers were estimated using the “elbow” method, where the number of clusters are weighed against the percent of variance explained by the clusters (31,32).

To validate the distinct composition of each cluster we first optimized a logistic regression model using backward elimination then trained the optimized model on the trauma cohort. We then validated on the two datasets with sepsis (abdominal and respiratory source). We also tested internal validation on the trauma dataset using k-fold cross-validation. Model performance was assessed using area under the receiver operator curve (33,34).

## Results

### Overlapping biological processes in critical care patients

Gene co-expression analysis, performed separately on each disease etiology with respective controls included, identified multiple expression modules which predominantly enrich in processes relating to inflammation and cell division (Figure 1).

**Figure 1:**
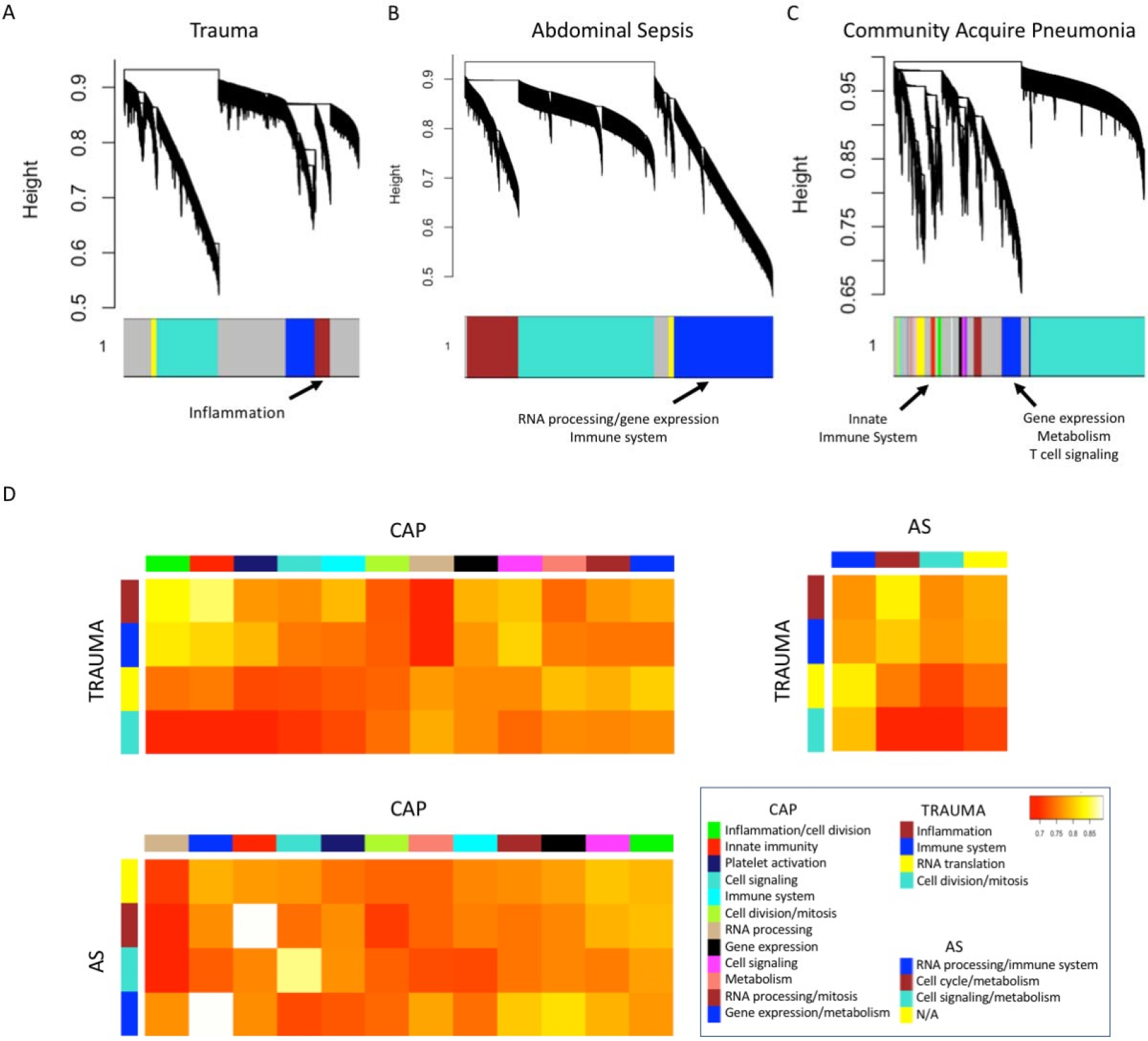
Shared transcriptome response across aetiologies of critical illness. Dendrogram summaries of gene co-expression networks derived from WBT in Trauma (A), Abdominal sepsis (B) and Community-acquired pneumonia (C). Color-coded bar graph under each dendrogram depicts co-expression modules, with grey representing unclassified genes. Only modules significantly different between disease and heathy controls are labeled (Mann Whitney p<0.05). Panel D shows heatmap summaries of ontological similarities between modules in sepsis vs trauma. Note the overlap for inflammation and immune processes across different aetiologies of critical illness. Panel *D bottom right*: GO terms enriching in each co-expression module. CAP: community acquired pneumonia, AS: Abdominal sepsis

Processes with the greatest degree of similarity between all disease types in GOSemSim analysis mapped largely to immune/inflammatory processes. This approach yielded a similarity metric greater than 0.75 between all modules which significantly differed between disease and controls.

### Distinct patterns of leukocyte compositions preserved across different aetiologies of critically illness

Motivated by the overlap in immunological and inflammatory processes we analyzed the immune cell composition from WBT of all 317 patients with critical illness. We applied the deconvolution technique CIBERSORT to compute relative fractions of 22 leukocyte subtypes from whole blood transcriptomics. This technique uses cell-specific marker transcripts to “deconvolve” lymphocyte cell fractions from bulk gene expression (12). Such deconvolution techniques are well established and have reshaped our understanding of the cell composition of “bulk tissue”. They have been applied in a number of studies to estimate not only leukocyte composition but also the immune landscape (18), microenvironment (19), and purity (35) of tumours. Deconvolution of leukocyte fractions showed complex but consistent patterns across trauma and sepsis caused by community acquired pneumonia or abdominal source (Figure 2A-C).

**Figure 2:**
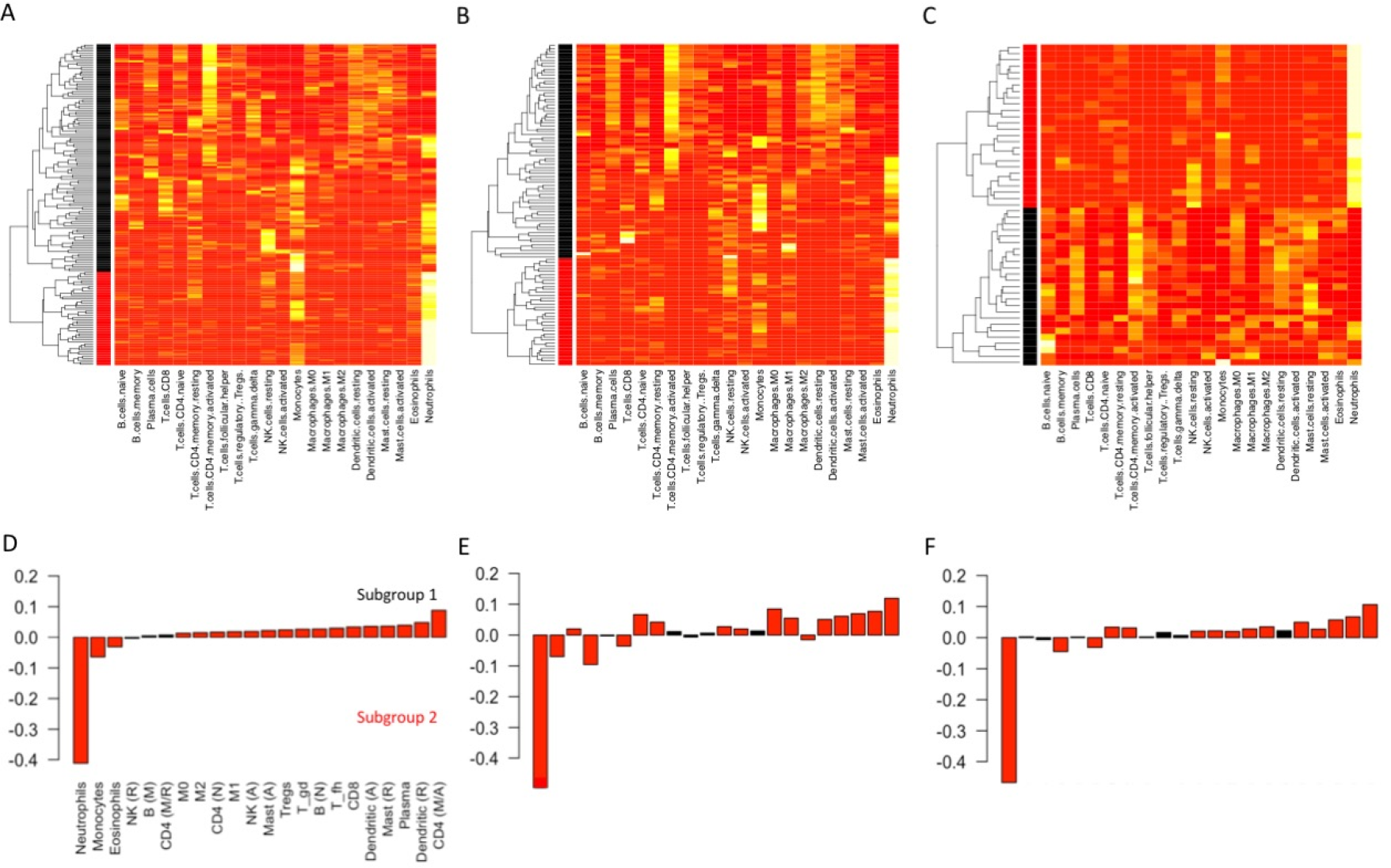
Shared immune cell configurations across aetiologies of critical illness. A-C: Heat map summaries of leukocyte composition in trauma (A), sepsis due to community acquired pneumonia (B) and sepsis due to an abdominal source (C). Sidebar indicates cluster assignment: black-subgroup 1, red-subgroup 2. Patients are presented in rows and cell types in columns. D-F: Bargraph summaries of differential leukocyte fractions showing suppressed neutrophils, elevated dendritic cells and CD4 memory T cells dominating subgroup 1 (top) and elevated neutrophils with otherwise suppressed leukocyte levels seen in subgroup 2 (bottom). Note the pattern remains consistent across the three independent aetiologies. Red bars indicate significant differences (t-test, p<0.05)

The broad characteristics of these distinct groups were as follows: subgroup 1: a higher fraction of neutrophils with relative suppression of leukocyte fractions, subgroup 2: a higher proportion of activated CD4 memory cells associated with resting/activated dendritic cells. We will be referring to these as subgroup 1 and subgroup 2 from here onwards. Comparative analysis of cell fraction for the two consistent subgroups is summarized in Figure 2D-F. Comparative analysis of patient subgroups did not reveal any differences in the clinical/demographics labels (age, gender or mortality) available in the data repositories (Table 1). We have further validated the distinct leukocyte composition of these two subgroups by training a logistic regression model on the trauma cohort, with subgroup as a binary response variable and cell fractions as regressors, to independently validate the model on the septic cohorts. Area under the ROC curve was over 0.98 for all groups, demonstrating that the two subgroups were highly distinct from one another and consistent in all 3 cohorts (Supplementary figure 1). For further verification, we demonstrated the generalizability of our results in a robust pediatrics dataset consisting of 181 patients with admitting diagnosis of sepsis from a variety of sources. Cluster detection using the same method as the discovery cohorts gave subgroup with similar characteristics: one group with CD4 helper/dendritic cell dominant (corresponding to subgroup 1) and a second neutrophil dominant corresponding to subgroup 2 (Supplementary figure 2).

### Suppressed immune and inflammatory mechanisms in distinct subgroup of critically ill

We finally analysed correlation between the patient subgroups and biological mechanisms in critical illness (Figure 3). This was done by correlating module meta genes (Figure 1) with group labels of individual patients (healthy, subgroup 1, subgroup 2). We found that modules that enrich in inflammatory/immune mechanisms were of highest significance. The expression of these modules (meta-genes) were higher in the subgroup 2 (neutrophil-dominated) and lower in subgroup 1 (CD4 T helpers/dendritic cell dominated). Analysis of our validation cohort paralleled these results: module meta genes from the pediatric cohort suffering from septic shock of diverse aetiologies correlated well with subgroup labels: subgroup 2 showed significantly higher correlation with innate immune mechanisms compared to subgroup 2 (Supplementary figure 3).

**Figure 3:**
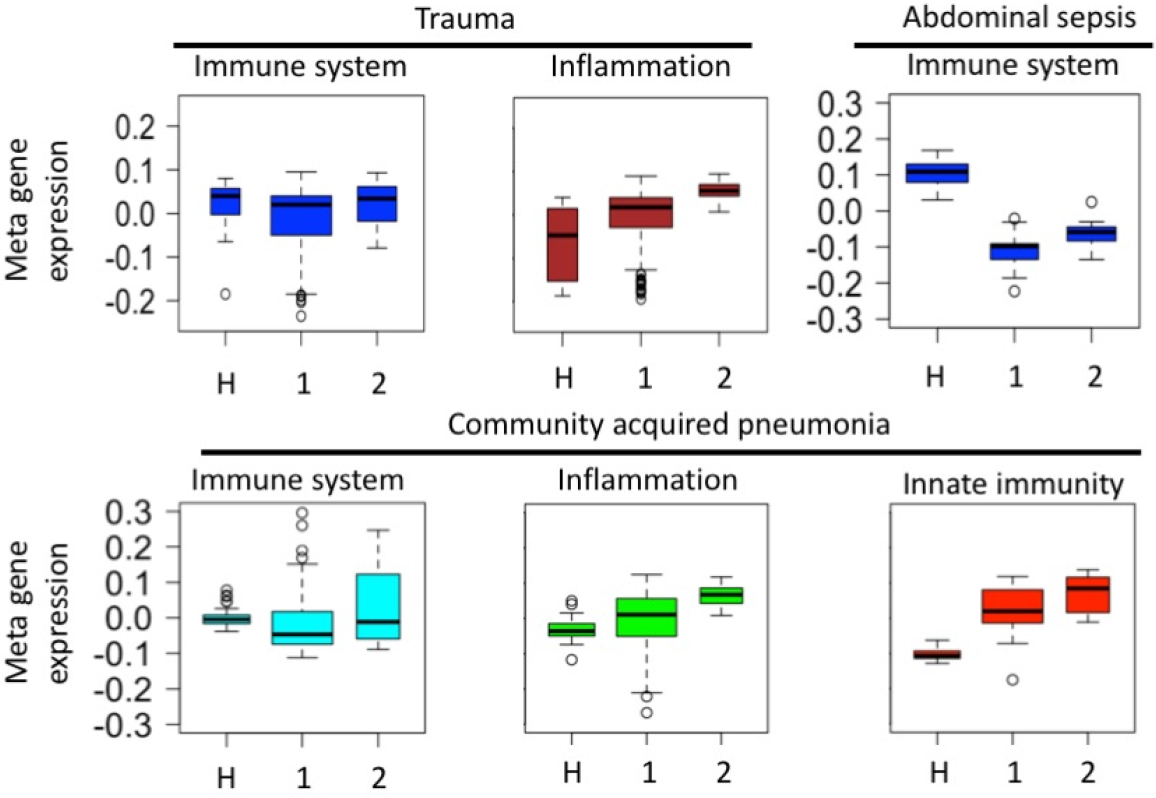
Boxplot summary of module meta-gene correlation with patient subgroup phenotypes: H: healthy, subgroup 1: CD4 T helper, dendritic cell predominant and neutrophil depleted, subgroup 2: Neutrophil predominant. Note the lower relative expression level of module eigengenes for subgroup 1 versus subgroup 2. All modules plotted as significantly different between subgroups 1 and 2 (MW p<0.05)

## Discussion

Our results are the first to show two groups of patients characterized by different patterns of immune response to severe community acquired pneumonia, abdominal sepsis and trauma. One consistent patient subgroup consists depleted neutrophils, high memory CD4 T and dendritic cells counts (subgroup 1) and a second is characterized by high neutrophil counts (subgroup 2). Our results are the first to show that patterns of leukocyte composition are preserved across these different critical illnesses and may give basis for a “pan-disease” marker for patients with a complicated clinical course. In addition, our study confirms the consistent but diverse activation of host immune mechanisms in both sepsis and trauma (1, 2–4). (1).

Analysis of serum transcriptomics in critical illness has demonstrated a large scale reprioritization of gene expression, also referred to as a “genomic storm”, following burns and trauma (2). Part of this effect is due to the up-regulation of innate immune mechanisms, microbial recognition and inflammation, and down-regulation of antigen presentation and adaptive immunity. Complicated recovery (ie: prolonged hospital stay, no recovery or death) was associated with upregulated cytokines IL-6 and IL-10, transcription factors p38, and MAPK signalling cascades, while antigen presentation and T-cell regulation is suppressed (2). The current models of the immune/inflammatory paradigm suggest two consecutive mechanisms for the initial surge of inflammatory processes (Systemic Inflammatory Response Syndrome) followed by a compensatory anti-inflammatory mechanism. These recent genome wide results have suggested a new model consisting of the parallel upregulation of innate immunity with the suppression of adaptive immunity (2). In a longitudinal analysis of gene expression levels and cell factions in the hyperacute phase of trauma showed a relatively limited upregulation of transcriptomics (3). Only 4.2%, 21.4% and 21.0% of total genes were differentially expressed in samples taken within 2 hours of injury, 24 and 72 hours of admission, respectively. Biological processes associated with Multi Organ Dysfunction Syndrome (MODS) were cell death and survival pathways rather than a pro-inflammatory state suggested by prior concepts. We used co-expression networks to detect gene modules in sepsis and trauma patients, yielding processes enriching in inflammation, innate immunity and cell division. These results are in line with previous findings wherein comparison of transcriptomic responses to fecal peritonitis and community acquired pneumonia showed 64% overlap in upregulated genes with trauma (6).

Multiple configurations in leukocyte fractions have been suggested to associate with increased morbidity/mortality in trauma and sepsis (36,37). These configurations were proposed to contribute to the dysregulated immune response and immune suppressed state seen in critical illness. Studies using flow cytometry have shown higher risk of MODS with leukopenia, increased NK dim cells and reduced **δγ** low T lymphocytes in septic patients (36). Analysis of activation markers CD11b and HLE for neutrophils, and ICAM-1 for monocytes showed reduced signals for patients with higher mortality in sepsis (37). Neutrophils have a central role in triggering and amplifying innate immunity. One of the subgroups identified in our study is dominated by a high fraction of neutrophils while other leukocyte fractions are suppressed. We were, however, limited to assessing cell fractions and lack information on the biological activity of these fractions. The second subgroup is characterized by elevated dendritic cells and memory CD4 T fractions. Memory CD4 T cells are derived from activated CD4 T cells (38); while their role is not well defined, they are proposed as early effectors of cytokine response, helpers of B/T cell response and effector responses as reviewed in ref 36 (39). However, it has been suggested that the apoptotic response in sepsis causes the recovery of an antigen specific cell pool with narrower repertoire, translating to a dysregulated immune response (40). Dendritic cells are regarded as the link between innate and adaptive immunity as they stimulate the adaptive immune system through their antigen presenting and secretory functions (41). In response to microbial stimuli, dendritic cells express co-stimulants such as CD40 and CD86 through a process termed maturation (42). The depletion of dendritic cells was shown to associate with increased frequency of ICU acquired infection for patients with septic shock (7) and in burns (43). Beyond just down-regulation, dendritic cells were proposed to contribute to immune-suppressed states in sepsis through alterations in their differentiation (44), phenotypical changes, and immunological functions (45). Furthermore, an important dichotomy in dendritic cell function is the secretion of IL-12 versus IL-10. IL-12 polarizes naïve Th to Th 1 promoting immune response by linking innate to adaptive immunity (46). Conversely, IL-10 promotes regulatory T cells and suppresses Th 1 priming, NKs and macrophages, hence promoting an immune-suppressed state (47). Given that our analysis is limited to cell fractions and lacks data on the functional state of these cells it is difficult to draw conclusions on the immune state of these subgroups. We were nevertheless able to correlate gene expression patterns that annotated to immunity and inflammation with the two subgroups in leukocyte configuration. We found downregulation of both immunity and inflammatory mechanisms in the first subgroup of depleted neutrophils, increased dendritic and CD4 helper cell fractions implying an immunosuppressed state. Supporting this interpretation are prior studies that correlate pure fractions of leukocytes to clinical course and outcome. Deconvolution of leukocyte fractions in trauma showed increased risk of Multi Organ Dysfunction Syndrome (MODS) with elevated dendritic cell counts but depleted neutrophil levels (3) which corresponds to the configuration of subgroup 1. Given the limitation of our dataset further studies are required to confirm the association of subgroup 2 with MODS in critical illness.

## Conclusion

Our study identifies two patient consistent subgroups in diverse forms of critical illness, distinguished based on their leukocyte composition. This preserved configuration may serve as a universal marker of multiple organ dysfunction for the critically ill, and allow for more targeted approaches to outcome prediction and treatment.

## Supplementary material

**Supplementary table 1:** Comparative analysis of patient endotypes distinguished based on immune cell compositions.

|  | Endotype | Mean age<br>(SD) | Sex<br>(M:F) | Mortality<br>(%) |
| --- | --- | --- | --- | --- |
| TRAUMA | Subgroup 1 | 33.4 (10.9) | 75:37 | 6/112 (5.4) |
|  | Subgroup 2 | 35.5 (11.9) | 26:20 | 1/46 (2.2) |
| ABDOMEN | Subgroup 1 | 62.2 (13.1) | 13:13 | 4/26 (15.4) |
|  | Subgroup 2 | 61.5 (10.9) | 14:11 | 4/25 (16.0) |
| CAP | Subgroup 1 | 60.6 (16.2) | 43:29 | 15/72 (20.8) |
|  | Subgroup 2 | 61.8 (17.2) | 20:16 | 8/36 (22.2) |

\*No parameters have $p < 0.05$ when comparing endotypes of the same cohort (t-test for age and chi square for sex and mortality).

**Supplementary figure 1:**
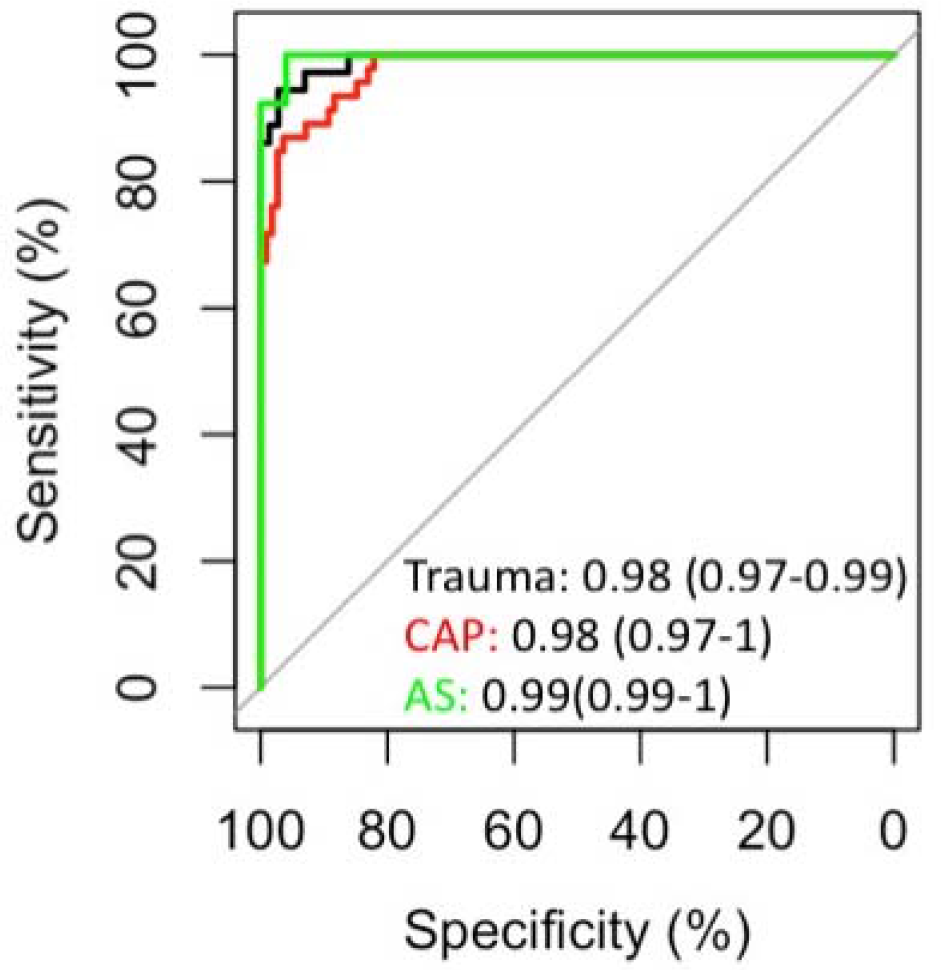
Validation of patient subgroups based on leukocyte composition. Area under (AUC) the receiver operator curve (ROC) show highly accurate classification of patient subgroups (AUC ≥ 0.98). AUC values are show as AUC (95% CI).

**Supplementary figure 2:**
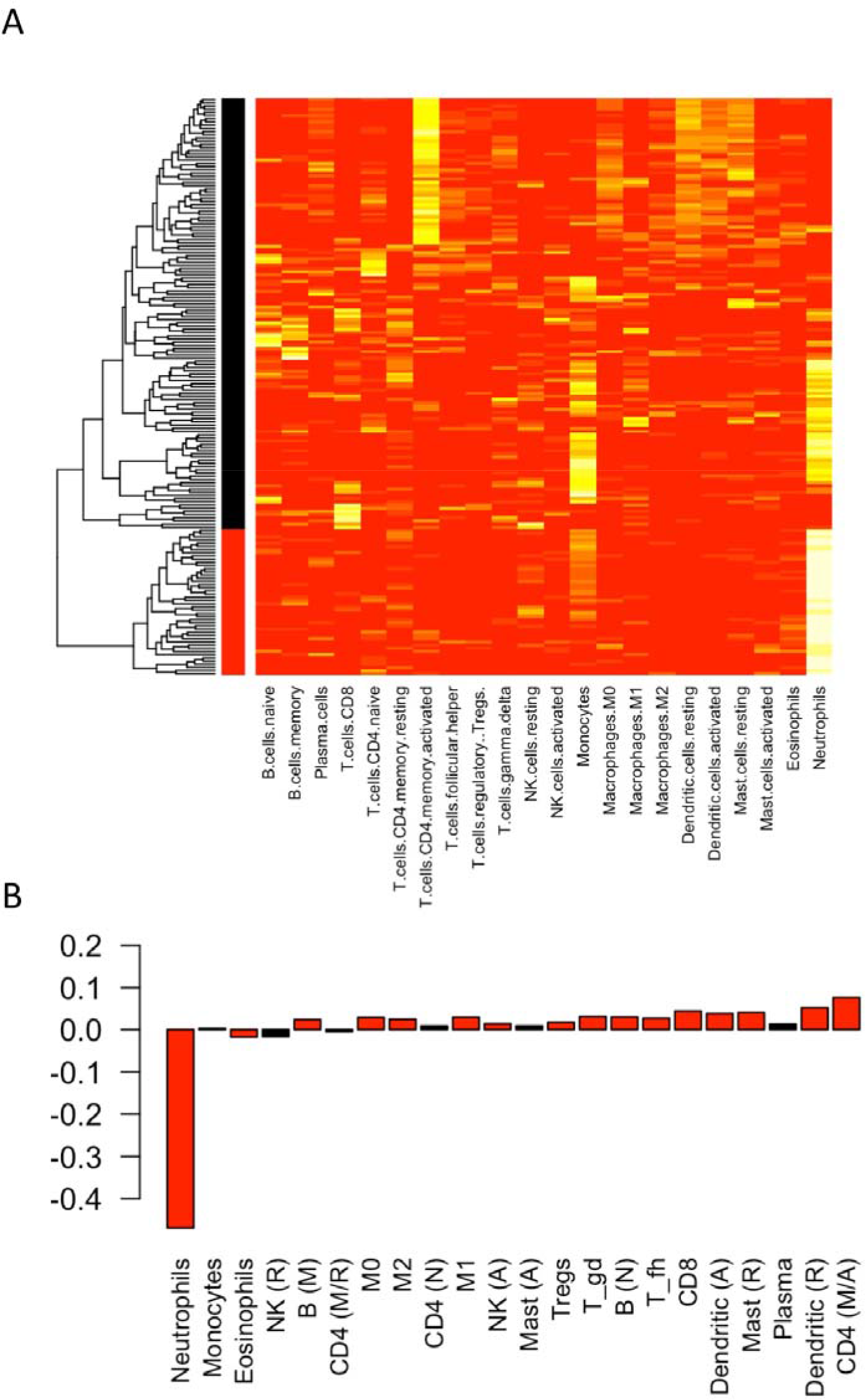
Verification of patient subgroups; one with increased neutrophils and second group with relatively supressed neutrophils and dominant dendritic cells, activated CD4 memory cells. A: Heatmap summary of clustering. Black: Subgroup 1, Red: Subgroup 2. B: Barplot representing differential fractions between subgroups 1 and 2. Red bars indicate significant difference between subgroups (MW p<0.05)

**Supplementary figure 3:**
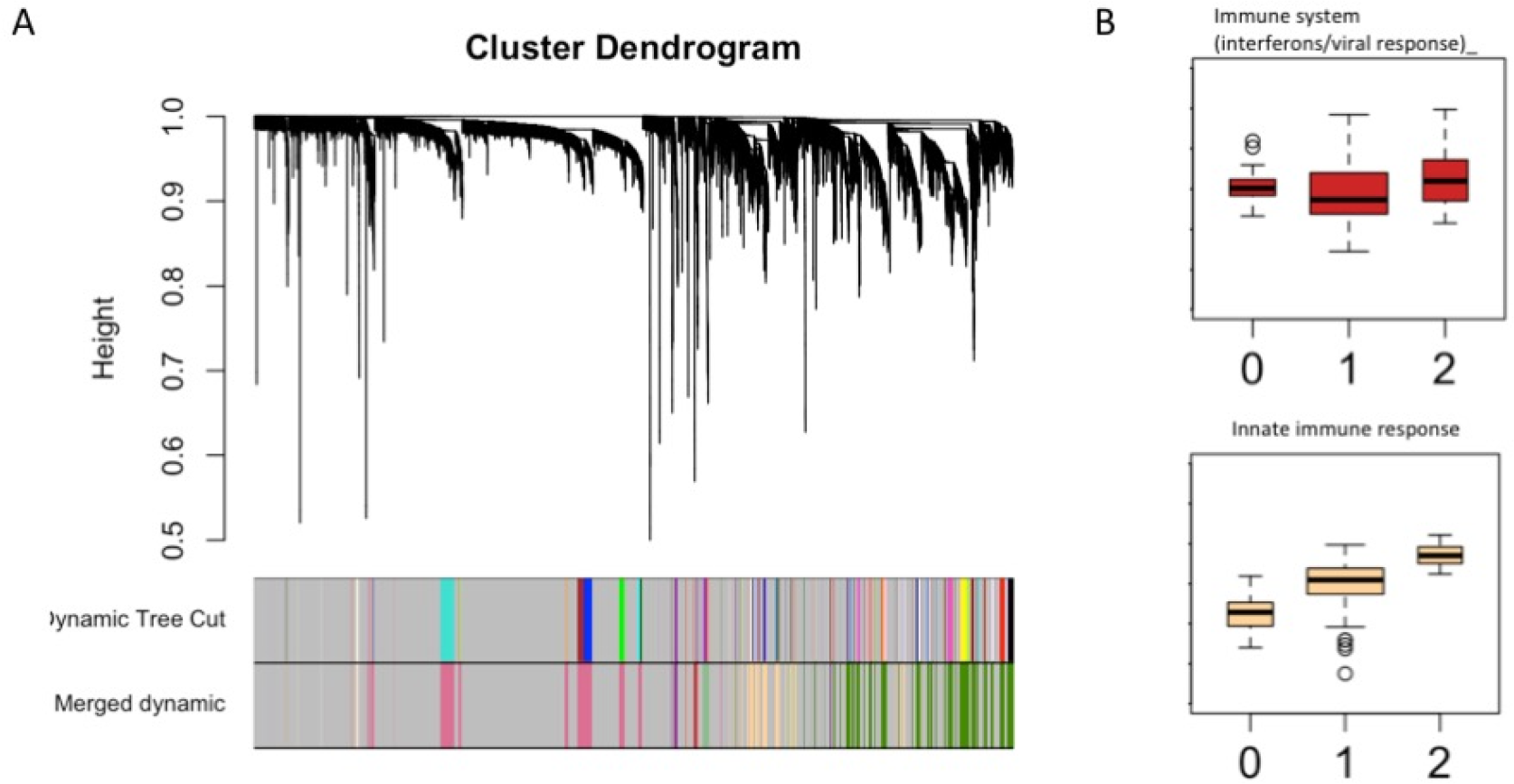
Verification of immune gene modules and meta gene correlations with patient subgroups in paediatric septic shock. Dendrograms with merged dynamic tree cut (A) gave gene co-expression modules with similar annotations as adult cohorts of trauma and sepsis. Panel B: Correlation of meta genes with subgroup memberships (Supplementary figure 2) paralleled results from adult cohort, further suggesting subgroup 2 with higher immune activity versus subgroup 1 corresponding to an immune supressed state. 0: healthy, 1: group 1 and 2: group 2

Author Contributions
ZZ: study design, data analysis, manuscript preparation
APL: data analysis, manuscript revisions
MB: data analysis, manuscript revisions
JCM: study design
MDC: manuscript revisions

